# An atlas of the catalytically active liver and spleen kinases in chicken identified by chemoproteomics

**DOI:** 10.1101/634683

**Authors:** Bindu Nanduri, Cathy R. Gresham, Winnie W. Hui, Mark Ou, Richard H. Bailey, Mariola J. Edelmann

**Author notes:** Corresponding author, Department of Microbiology and Cell Science, University of Florida, Gainesville, FL, 32611.

## Abstract

Chicken is the first agricultural animal to have a sequenced genome, but current kinase annotations of *Gallus gallus* are overwhelmingly limited to the predictions generated based on homology or isolated studies focused on specific kinases. Our approach used chemical probes consisting of ATP and ADP derivatives binding to specific lysine residues within the ATP-binding pocket of kinases. Collectively, we identified 188 chicken kinases and corresponding 267 peptides labeled with the ATP and ADP acyl derivatives in chicken spleen and liver. The kinases identified here are publicly available through the database, Chickspress genome browser (*http://geneatlas.arl.arizona.edu/cgi-bin/GeneAtlas/chickspress_genes.cgii*). Analysis of putative functions of these chicken kinases indicates that kinases identified in this study might regulate hematological system development, necrosis, apoptosis, epithelial neoplasm, and other processes. The identified tissue-specific expression atlas of active chicken kinases along with the ATP binding sites of kinases provide the basis for the development of specific drug targets for multiple chicken diseases as well as starting point for inhibitor selectivity studies in this agriculturally important species. Moreover, this study will support future studies focused on identifying the role of these kinases in chicken growth, metabolism, and disease.

## Introduction

Chicken represents an essential source of protein. The U.S. poultry industry is the largest in the world and continues to be a major supplier of meat and eggs. The world’s population growth is projected to surpass 9 billion by 2050, which mandates intensification of the existing poultry operations. It is necessary to make improvements in the breeding, management, and treatment of poultry to meet this projected higher demand for food. A comprehensive understanding of the physiology and pathophysiology of the chicken is critical for animal production. Chicken is the first agricultural animal to have a sequenced genome (1), with a description of the protein-coding genes in the genome. Understanding the biological role of annotated gene products, i.e., proteins require a description of their biological function. Like other non-model agricultural organisms, the current annotation of the protein-coding genes in the chicken genome relies on computational predictions based on sequence similarity, based on the ortholog conjecture that postulates functional similarity of orthologous genes (2, 3). The dynamic response, regulation, and modification of the genome ultimately governs complex phenotypes. The availability of the chicken genome expedites knowledge discovery using genomics methodologies. A compendium of omics approaches such as transcriptomics, proteomics, or metabolomics enables the measurement of the genome-scale response of chicken at the RNA, protein, and metabolite levels, respectively, to identify molecular mechanisms responsive to biotic and abiotic perturbations. Several post-transcriptional regulatory mechanisms such as protein post-translational modifications (PTMs) including, but not limited to, phosphorylation, glycosylation, ubiquitination, and methylation are known to regulate protein structure and function. Post-translational regulatory mechanisms govern quick response to external/internal stimuli and changes in PTM are often associated with the disease. Protein phosphorylation is the most common and best characterized among PTMs, and it is estimated to affect approximately 30% of all proteins in eukaryotes (4, 5). Protein kinases catalyze the enzymatic addition of phosphoryl groups from adenosine triphosphate (ATP) to the hydroxyls of specific amino acid residues, while phosphatases reverse this modification. Phosphorylation of proteins catalyzed by kinases occupies a pivotal role in signal transduction pathways in eukaryotes and is known to control a wide range of cellular processes, ranging from metabolism, proliferation, cell cycle progression, apoptosis and pathogen clearance (6, 7) (8). Protein phosphorylation is integral to organismal development, homeostasis, immune and nervous systems, and dysregulation. Over 500 kinases constitute the full complement of protein kinases in the human genome, also known as kinome. Protein kinases play regulatory roles in cancer, inflammatory disorders or infections (6, 9–15). Since protein kinases are associated with pathophysiology, they are attractive drug targets and are indeed among the most highly targeted class of enzymes in drug development. At present, there are 37 kinase inhibitors licensed for therapeutic use by US Food and Drug Administration ^11, 12, 16^ while 250 inhibitors are under clinical evaluation (16). Some of these drugs, such as tyrosine kinase inhibitors, are also explored in veterinary medicine applications (17). There are also efforts to repurpose kinase inhibitors as antimicrobials (18), and such novel classes of antibiotics could find use in treating multidrug-resistant bacteria, which affect poultry (19–21). Global characterization of chicken kinome will facilitate systems approaches to understand health and disease for developing novel therapeutics targeting this druggable class of enzymes. Species-specific peptide arrays can be utilized for studying kinomes, and peptide arrays that can interrogate cellular processes such as innate and adaptive immunity, inflammation, or protein, fat, and carbohydrate metabolism are described in the literature for poultry (22, 23). The available poultry-specific peptide arrays will help study the role of specific kinases in immune-metabolism. Mass spectrometry-based chemoproteomics approach utilizing probes that target ATP binding sites in catalytically active kinases is a complementary approach to identify a large number of active kinases in a given sample including potentially novel kinases missed by computational prediction. Chemoproteomics can help generate a tissue atlas for active kinases in an organism. A description of the full complement of catalytically active kinases in different chicken tissues will contribute to the current knowledge of kinome and provide information regarding the expression and activity of kinases which will aid future drug development in multiple domains such as physiology, development, immunology and host-pathogen interactions. For instance, activity screening by chemoproteomics is used for drug selectivity studies (24). Inhibitor selectivity can also be detected by other methods, such as competition experiments, which however require the use of recombinantly expressed kinase domains. Several chicken kinases are already known to regulate key cellular processes (25, 26). Expanding the current knowledge of the chicken kinome will further aid ongoing, and future studies to understand the interplay of kinases in different biological responses for the discovery of novel therapeutic interventions, such as selectivity studies of kinase inhibitors in chicken.

In this study, we utilize a chemoproteomics approach to identify active protein kinases in chicken liver and spleen and thus interrogate chicken kinome in these tissues. Our approach used chemical probes consisting of ATP and ADP derivatives binding to specific lysine residues within the ATP-binding pocket of kinases. Collectively, we identified 188 chicken kinases and corresponding 267 peptides labeled with the ATP and ADP acyl derivatives in chicken spleen and liver. Analysis of putative functions of these kinases indicates that active kinases identified in this study are known to regulate hematological system development, necrosis, apoptosis, epithelial neoplasm, and other processes. The identified chicken kinome along with the ATP binding sites of kinases provide the basis for the development of specific drug targets for multiple chicken diseases as well as starting point for inhibitor selectivity studies in this agriculturally important species. Kinases identified in this study are available at the Chickpress Genome Browser (http://geneatlas.arl.arizona.edu/cgi-bin/GeneAtlas/chickspress_genes.cgi)

## Results

### Chemoproteomics-based identification of protein kinases in chicken liver and spleen

We utilized a chemoproteomics approach for genome-wide identification of active kinases in chicken tissues. In this strategy, biotinylated acyl phosphates of ATP and ADP were used to label catalytically active lysine-containing domains of protein kinases present in liver and spleen homogenates. These acyl phosphate derivatives irreversibly bind to the conserved lysine residues present within the ATP-binding pockets of kinases (27). Protein kinases labeled with biotinylated probes were identified by mass spectrometry-based proteomics. We identified 267 peptides labeled with acyl derivatives of ATP and ADP that correspond to 188 proteins with kinase activity. We identified 183 protein kinases from liver and 188 kinases in the spleen. The kinases identified in this project are available for broader dissemination to the research community through the Chickpress Genome Browser (*http://geneatlas.arl.arizona.edu/cgi-bin/GeneAtlas/chickspress_genes.cgi*). A description of the identified ATP-binding site and abundance in tested tissues for all protein kinases reported in this study is given in **Supplementary Table 1**, and displayed as dendrograms, where the height of the bar corresponds to the MS signal strength, which correlates with the activity/abundance of the proteins (**Fig. 1, 2**). The ATP/ADP probes bind kinases with available ATP/ADP binding sites and identify kinases that are activated under the conditions of sample collection. Although results of this approach and other similar experimental approaches for identifying active kinases need to be further supported by other activity assays for individual kinases, inhibition data from KiNativ probes enriching kinases has a better correlation with readouts of cellular inhibition compared with other tests measuring kinase activity (24). Our results show that the spleen contained a higher number of kinases with available ATP/ADP binding sites in comparison to the liver (4.8-and 7-fold higher for ATP-and ADP-probes, respectively, when spectra of all identified active peptides were summarized for each probe **Supplementary Table 1**). The ATP/ ADP probes targeted a single site, and in some cases, more than one site **(Supplementary Table 1**). MET kinase is the only kinase which was relatively more abundant in the liver (∼5.5 fold) compared to the spleen. MET is critical in the development and regeneration of liver (28).

**Figure 1:**
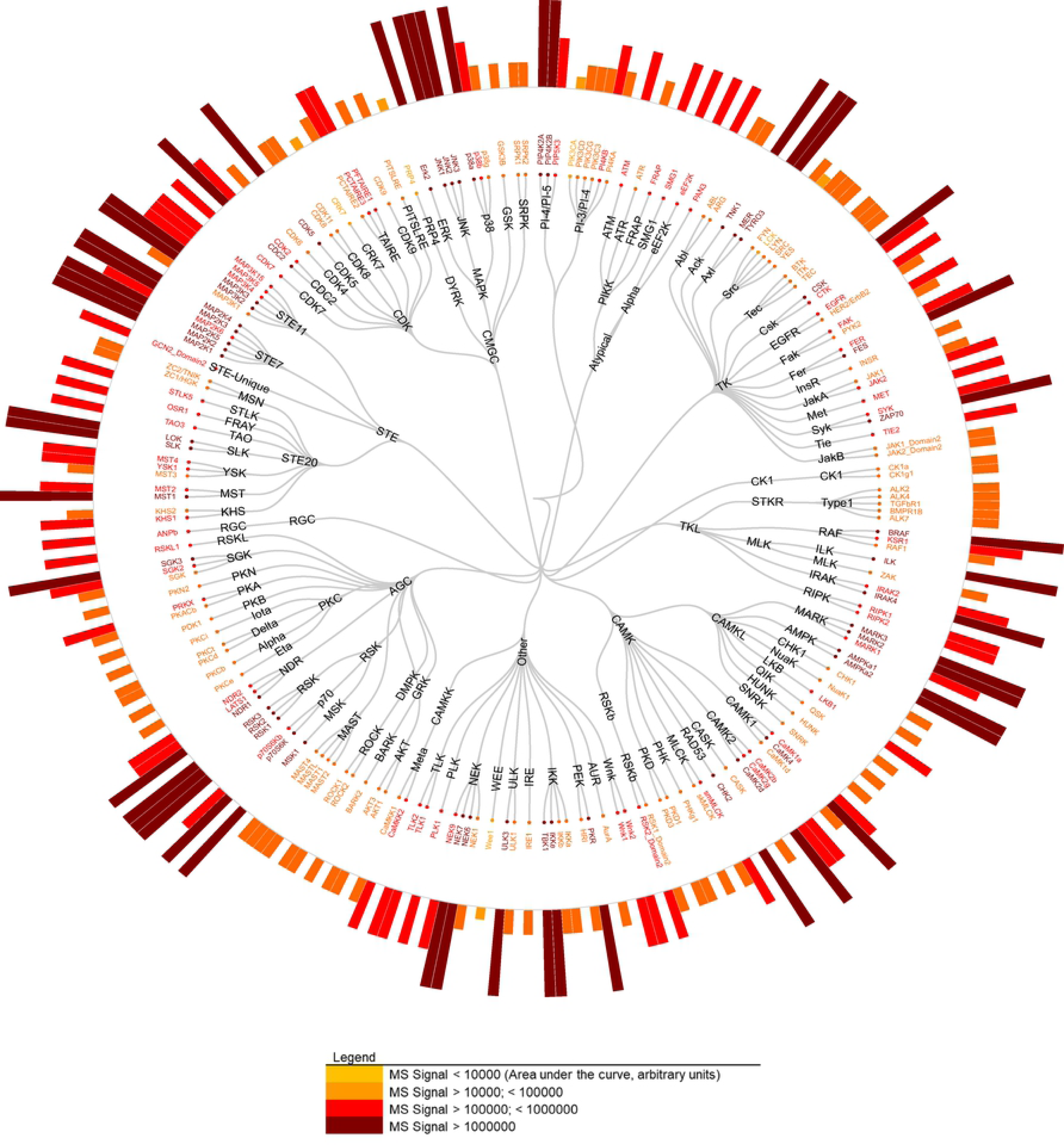
A dendrogram representing the identified chicken liver kinome. The protein kinases identified in the chicken liver through a chemoproteomics approach are organized into kinase groups followed by families (from top to bottom). The bars indicate the level of abundance of active sites, based on the mass spectrometry signal where maroon represents the most abundant kinases compared to yellow which represents the lowest abundance. Each kinase is color-coded for the peptide that yielded the highest signal.

**Figure 2:**
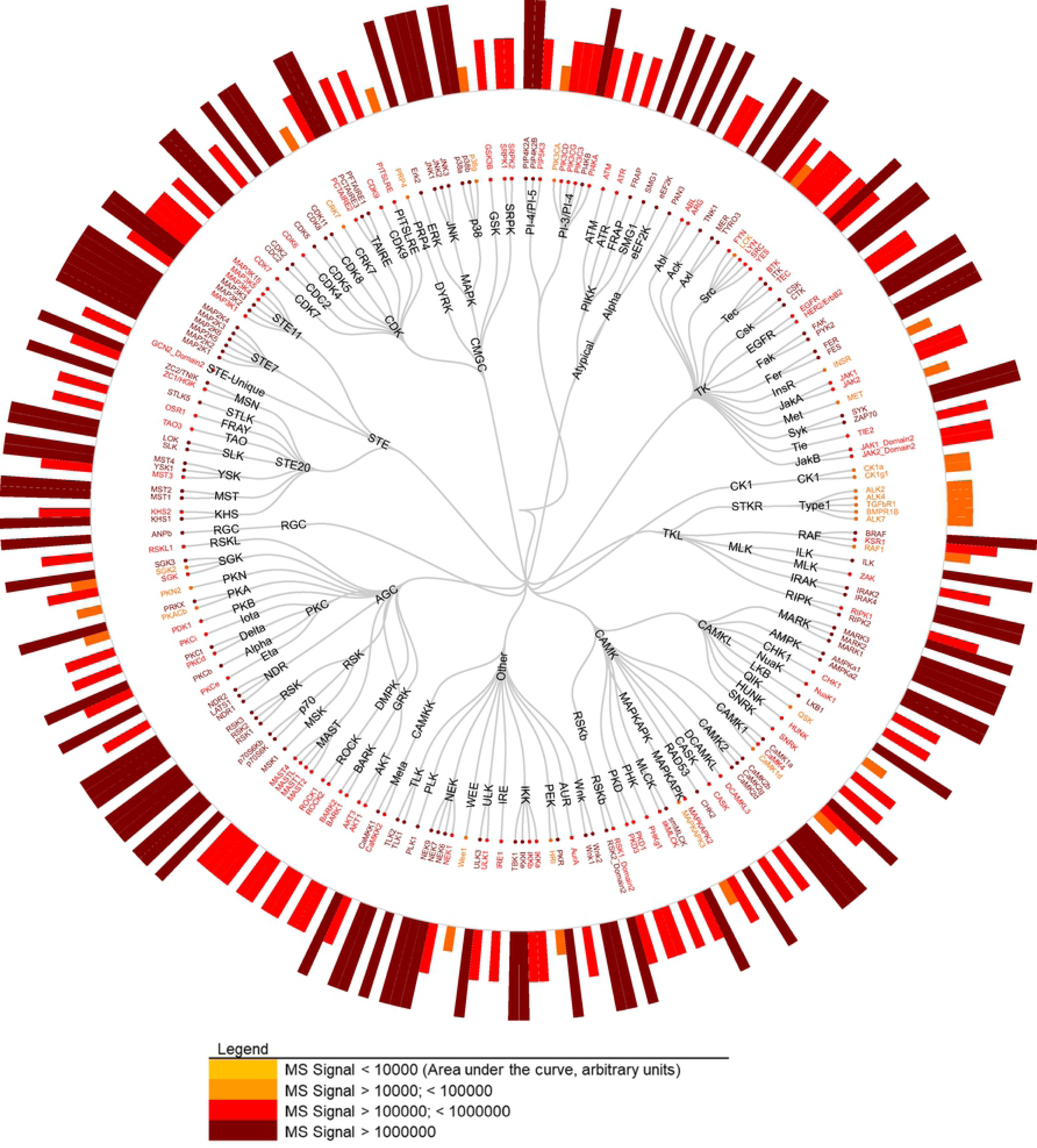
A dendrogram representing the identified chicken spleen kinome. The protein kinases, identified in the chicken spleen by a chemoproteomics approach, are organized into kinase groups followed by families (from top to bottom). The bars indicate the level of abundance of active sites, based on the mass spectrometry signal where maroon represents the most abundant kinases compared to yellow which represents the lowest abundance. Each kinase is color-coded for the peptide that yielded the highest signal.

The results of chemoproteomics-based identification of active kinases were validated via immunoblotting for three kinases, IKKa, AKT1, and PIK3CA. Western blotting confirmed higher expression of all three kinases in spleen compared to the liver (**Fig. 3A, B**).

**Figure 3.**
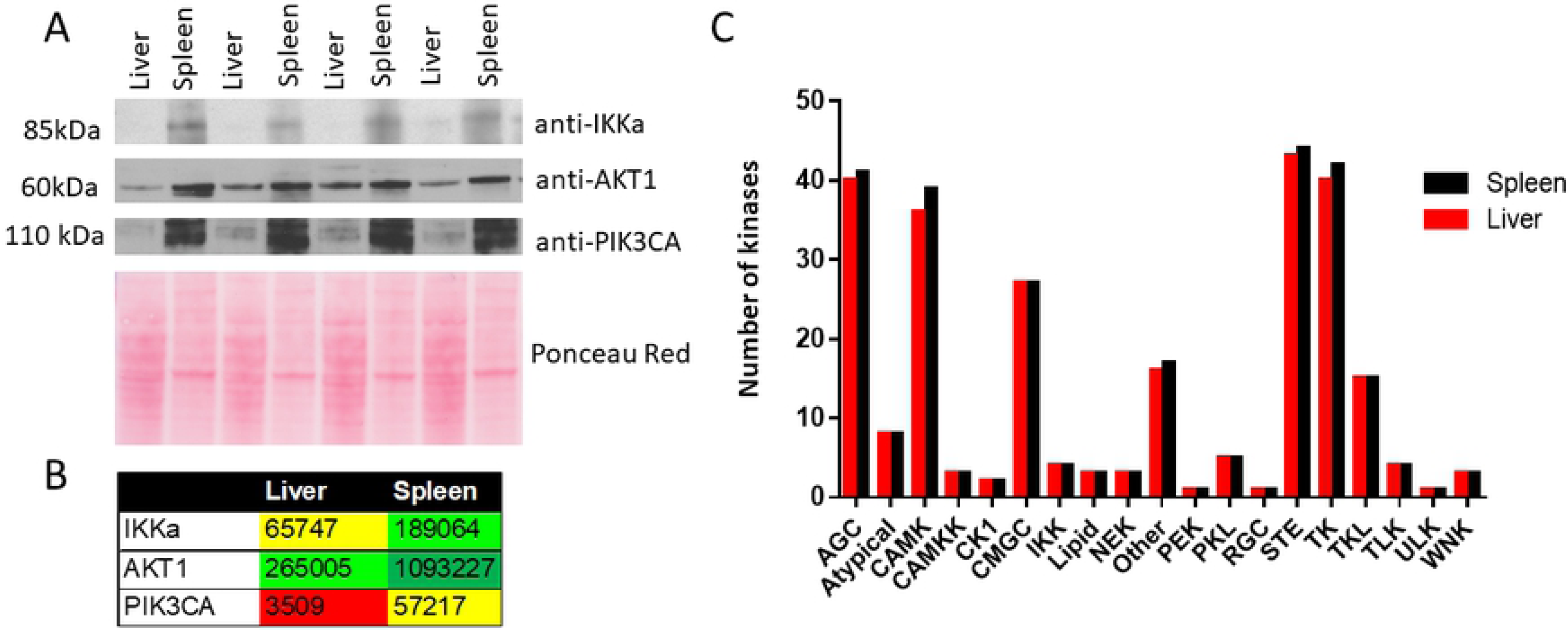
Detection of kinases by Western blot immunoassay. **(A)** An equal amount of protein from three liver or spleen samples (each tissue was harvested from a different bird) were separated by SDS-PAGE, followed by transfer onto PVDF membrane. The membrane was probed for IKKa, AKT1 or PIK3CA kinases, or stained with Ponceau Red stain to show protein loading. **(B).** Spectral counts for peptides of kinases identified by chemoproteomics (**Supplementary Table 1**) are shown to compare the intensities of proteins with mass spectrometry data.

The distribution of kinase families and subfamilies represented by the protein kinases identified in this study are shown as dendrograms (**Fig. 1, 2**). We identified a number of kinases in each of the nineteen individual most represented families in the liver and spleen (**Fig. 3C**).

Major kinase families identified in both liver and spleen include AGC, CMGC, CAMK, CK1, Other, STE, Tyrosine Kinase (TK), Tyrosine Kinase-Like (TKL), RGC, and Atypical kinases (**Fig. 1, 2, 3**). The AGC kinase group contains essential kinase families (29, 30), such as protein kinase A, G, and C, which are regulated by secondary messengers including cAMP, cGMP, or lipid molecules. In the human genome, there are 63 AGC kinases, and we identified 32 members of this kinase group in chicken (in equal number in spleen and liver), which is approximately half of the known AGC kinases in human. Another identified kinase group, CMGC, is named after CDK, MAPK, GSK3 and CLK kinase families. CMGC kinases include functionally diverse enzymes, which control the cell cycle, MAPK signaling, transcription, and cellular communication. In the human kinome, there are 57 CMGC kinases (31) of which 24 are identified in the chicken kinome reported here. STE kinases include the MAPK cascade families that constitute critical kinases for transducing signals, such as the highly abundant MAP2K and MAP3K kinases in the liver and spleen. While there are 47 known STE kinases (6) in human, we identified 27 chicken STE kinases, which constitute over 57% of the human STE kinases. Based on these examples, our study identified a substantial number of kinases from different protein kinase groups in chicken. However, our data also failed to identify 40-50% of the chicken orthologs of human kinases. This result indicates that at least some of these kinases are probably not active under experimental conditions, and some could be unique to human kinome, or their abundance was too low for confident detection in chicken tissues.

Analysis of the position of the labeled lysine residue within specific kinase domains showed that107 sites were mapped to the Lys1 domain and 91 to the Lys2 domain. Lys1, Conserved Lysine 1, is a residue localized at the beginning of the kinase domain, while Lys2, Conserved Lysine 2, is located in the middle of the protein kinase domain. In addition, 34 identified labeled sites were within the activation loop of the kinase domain, 21 were mapped to the ATP site in non-canonical kinase (**Figs. 1-2** “ATP”), and seven to ATP-binding loop. Finally, four sites were within other protein kinase domain (possibly not in the ATP binding site), while three others were not categorized, as the labeling of the residue was likely outside of the protein kinase domain and possibly not in ATP binding site.

### Conservation of kinase active site between chicken and other species

Sequence alignments were performed to determine the level of conservation between the identified peptides of chicken kinases and their closest human, rat and mouse homologs. The known ATP-/ ADP-binding sites of human and mouse proteins were aligned with the regions of the identified *Gallus* proteins, including the ATP-binding lysine residues, which is labeled by the biotinylated acyl phosphates and detected by mass spectrometry. A protein-BLAST with three chicken serine/threonine-protein kinases, WNK1, SMG1, and Nek6 identified in this study was performed to identify human, rat and mouse homologs. Protein sequences of chicken, human, rat, and mouse kinases were aligned and visualized (**Fig. 4**). The ATP-binding lysine residues of WNK1 and Nek6 kinases in mouse, rat and human proteins (shown as a blue box) corresponded to a lysine residue in chicken kinase (shown as a red box). The lysine residue in chicken kinase was modified by the ATP-or ADP-acyl phosphate probes in this study. The identified ATP-binding residue in chicken kinase SMG1 has not been annotated as an ATP binding site in human, rat and mouse SMG1.

**Figure 4:**
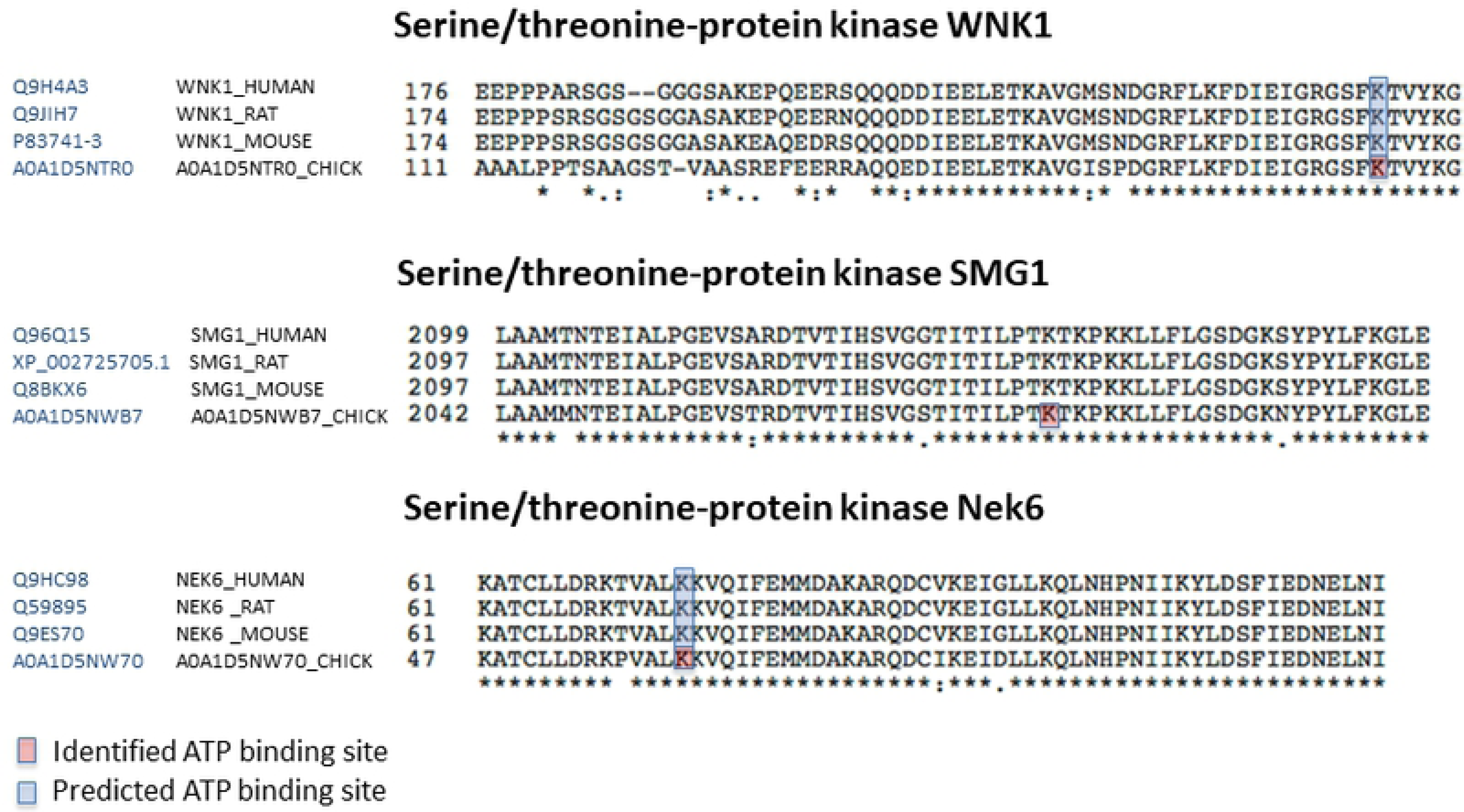
Sequence alignments of representative chicken kinases identified through chemoproteomics. Three *Gallus gallus* proteins were chosen for sequence alignment. We used *blastp* to identify the reviewed orthologs of chicken kinases in human, rat, and mouse. Red boxes indicate the residue, which was identified by mass spectrometry to bind to the active-site biotinylated acyl phosphate probe (**Supplementary Table 1**) and therefore predicted to bind ATP/ADP. The blue boxes indicate known ATP-binding regions in orthologous proteins in other species.

### Analysis of the biological functions of chicken liver and spleen kinases

To determine the biological function of the identified kinases reported here, we performed Ingenuity Pathway Analysis. Based on the annotated functions of human, rat, and mouse kinases, chicken kinases identified in the spleen (**Fig. 5A**) and liver (**Fig. 5B**) were associated with tissue development, hematological system development, lymphoid tissue structure and development, organismal survival and hematopoiesis. Chicken kinases identified by chemoproteomics also were associated with disease etiology (**Table 1-2**). Kinases identified in the liver were mapped to specific liver conditions such as a lesion, apoptosis of hepatoma cells, necrosis of the liver, liver tumor, differentiation of erythroid precursor cells, epithelial neoplasm and others (**Table 1**). Kinases identified in chicken spleen were mapped to critical functions of spleen including morphology of spleen and lymphoid tissue, the quantity of blood cells, the proliferation of lymphatic system cells, cell death of immune cells, the viability of B lymphocytes and others (**Table 2**).

**Figure 5.**
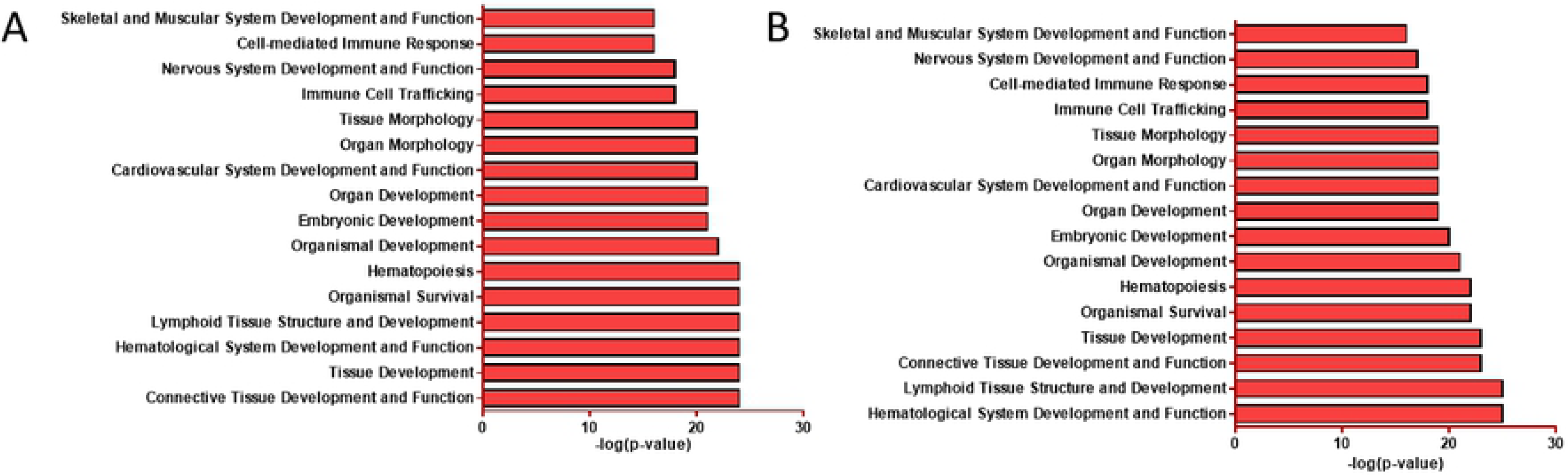
Physiological system development and functions associated with kinases identified by Ingenuity Pathways Analysis. All kinases identified in the spleen (A) or liver (B) tissue by Mass Spectrometry were analyzed by IPA to determine the physiological functions associated with these proteins.

**Table 1.** **Biological functions and hepatic diseases associated with chicken liver kinases identified** by Ingenuity Pathway Analysis.

| Biological function/disease | p-Value | Kinases | total # of kinases |
| --- | --- | --- | --- |
| Cell death of hepatoma cell lines | 3.3E-08 | AKT1,EGFR,ERN1,INSR,IRAK2,MAP2K1,MAP2K3,MAP2K4,MAP3K7,MAPK14,MET,PLK1,PRKCD,PRKCZ,SYK,ULK1 | 16 |
| Apoptosis of hepatoma cell lines | 4.3E-07 | AKT1,EGFR,INSR,MAP2K1,MAP2K3,MAP2K4,MAP3K7,MAPK14,MET,PLK1,PRKCD,PRKCZ,SYK,ULK1 | 14 |
| Liver lesion | 2.4E-05 | ABL1,ATM,ATR,BTK,CAMK2B,CAMK2D,CAMK2G,CDK2,CDK9,CHUK,EGFR,EIF2AK1,INSR,ITK,JAK1,LATS1,MAP2K1,MAP2K3,MAP2K4,MAP3K1,MAP3K4,MAP3K7,MAPK14,MAPK9,MAST2,MAST4,MERTK,MET,MTOR,MYLK,NEK6,NPR2,PAN3,PDPK1,PI4KA,PIK3CA,PKN2,PRKAA2,PRKCB,PRKCQ,PRKD3,RAF1,RIPK1,RIPK2,ROCK1,RPS6KA2,RPS6KA5,SGK1,SGK2,SIK2,SLK,SNRK,SRPK1,STK11,STK25,STK4,SYK,TAOK3,TBK1,TNIK,TNK2,WEE1 | 62 |
| Cell death of liver cells | 2.9E-05 | CHUK,EGFR,MAP2K1,MAP2K4,MAPK14,MAPK9,MET,MTOR,PRKAA2,RAF1,RIPK1,STK4,TBK1 | 13 |
| Necrosis of liver | 4.1E-05 | ABL1,CHUK,EGFR,MAP2K1,MAP2K4,MAPK14,MAPK9,MET,MTOR,PRKAA2,RAF1,RIPK1,STK4,TBK1 | 14 |
| Liver tumor | 0.00013 | ABL1,ATM,ATR,BTK,CAMK2B,CAMK2D,CAMK2G,CDK2,CDK9,EIF2AK1,INSR,ITK,JAK1,LATS1,MAP2K3,MAP2K4,MAP3K1,MAP3K4,MAP3K7,MAPK14,MAPK9,MAST2,MAST4,MERTK,MET,MTOR,MYLK,NEK6,NPR2,PAN3,PDPK1,PI4KA,PIK3CA,PKN2,PRKAA2,PRKCB,PRKCQ,PRKD3,RAF1,RIPK1,RIPK2,ROCK1,RPS6KA2,RPS6KA5,SGK1,SGK2,SIK2,SLK,SNRK,SRPK1,STK11,SYK,TAOK3,TBK1,TNIK,TNK2,WEE1 | 57 |
| Necrosis of epithelial tissue | 0.00015 | ABL1,CHUK,EGFR,MAP2K1,MAP2K4,MAPK9,MET,MTOR,PRKAA2,RIPK1,STK4 | 11 |
| Differentiation of erythroid precursor cells | 0.00022 | ABL1,EGFR,EIF2AK1,JAK2 | 4 |
| Proliferation of hepatocytes | 0.00033 | AKT1,CDK2,EGFR,INSR,MAPK1,MAPK14,MAPK9,MET,RAF1,RPS6KB1 | 10 |
| Epithelial neoplasm | 0.00033 | ABL1,ATM,ATR,BTK,CAMK2B,CAMK2D,CAMK2G,CDK2,CDK9,EIF2AK1,INSR,ITK,JAK1,LATS1,MAP2K3,MAP2K4,MAP3K1,MAP3K4,MAP3K7,MAPK9,MAST2,MAST4,MERTK,MET,MTOR,MYLK,NEK6,NPR2,PAN3,PI4KA,PIK3CA,PKN2,PRKAA2,PRKCB,PRKCQ,PRKD3,RAF1,RIPK1,RIPK2,ROCK1,RPS6KA2,RPS6KA5,SGK1,SGK2,SIK2,SLK,SNRK,SRPK1,STK11,SYK,TAOK3,TBK1,TNIK,TNK2,WEE1 | 55 |
| Apoptosis of hepatocytes | 0.00061 | CHUK,EGFR,MAP2K1,MAP2K4,MAPK9,MET,MTOR,RIPK1,STK4 | 9 |
| Cell viability of hepatoma cell lines | 0.0007 | AKT1,MET,MTOR,PRKAA2,PRKCD | 5 |
| **Viral Infection** | 0.001 | AKT1,CAMK1,EGFR,EIF2AK2,INSR,PI4KA,RAF1,SRPK1 | 8 |

**Table 2.** Biological functions and diseases associated with chicken spleen kinases identified by Ingenuity Pathways Analysis.

| **Biological function/disease** | **p-Value** | **Kinases** | **total # of kinases** |
| --- | --- | --- | --- |
| **Morphology of lymphoid tissue** | 2E-05 | ABL1,ATM,AURKA,BTK,FYN,JAK2,LYN,MAPK14,MAPKAPK2,MERTK,MTOR,PDPK1,PIK3CD,PRKCD,STK11,STK4,TBK1,TYRO3,ULK1 | 19 |
| **Quantity of lymphatic system cells** | 3.1E-05 | ATM,AURKA,BTK,CAMK2G,CAMK4,FYN,LYN,MAPK14,MAPK9,MTOR,PIK3CD,PIK3CG,PRKCB,PRKCQ,RIPK2,STK11,STK4 | 17 |
| **Morphology of spleen** | 3.5E-05 | ABL1,ATM,AURKA,BTK,FYN,JAK2,LYN,MAPK14,MERTK,MTOR,PDPK1,PIK3CD,PRKCD,STK11,TBK1,TYRO3,ULK1 | 17 |
| **Quantity of blood cells** | 3.5E-05 | AURKA,BTK,CAMK2G,CAMK4,FYN,JAK2,LYN,MAPK14,MAPK9,MTOR,PIK3CD,PIK3CG,PRKCB,PRKCQ,RIPK2,STK11,STK4 | 17 |
| **Cell viability of lymphocytes** | 4.6E-05 | AKT1,BTK,CHEK2,PIK3CA,PIK3CD,PIK3CG,PRKCQ,SYK | 8 |
| **Mobilization of Ca<sup>2+</sup>** | 4.6E-05 | AKT1,BTK,PIK3CA,PIK3CD,PIK3CG | 5 |
| **Proliferation of lymphatic system cells** | 4.6E-05 | ABL1,AKT1,ATM,BTK,FYN,ITK,JAK2,LYN,MERTK,MTOR,PIK3CA,PIK3CD,PIK3CG,PRKCQ,SYK | 15 |
| **Cell death of immune cells** | 0.00012 | AKT1,ATM,BTK,CHEK2,ITK,PIK3CA,PIK3CD,PIK3CG,PRKCQ,SYK | 10 |
| **Cell viability of B lymphocytes** | 0.00015 | AKT1,BTK,PIK3CA,PIK3CD,PIK3CG,SYK | 6 |
| **Proliferation of B lymphocytes** | 0.00037 | ABL1,AKT1,MERTK,MTOR,PIK3CA,PIK3CD,PIK3CG,SYK | 8 |
| **Proliferation of lymphocytes** | 0.00039 | ABL1,AKT1,ATM,FYN,ITK,MERTK,MTOR,PIK3CA,PIK3CD,PIK3CG,PRKCQ,SYK | 12 |
| **Activation of cells** | 0.00045 | CHUK,ITK,LYN,MERTK,PTK2B,RIPK2,STK11,SYK | 8 |
| **Activation of B lymphocytes** | 0.00053 | LYN,MERTK,RIPK2,SYK | 4 |
| **Expansion of CD4<sup>+</sup> T-lymphocytes** | 0.00078 | PIK3CA,PIK3CD,PIK3CG | 3 |
| **Quantity of T lymphocytes** | 0.00094 | BTK,CAMK2G,CAMK4,FYN,MAPK14,MAPK9,PIK3CG,PRKCQ,STK11,STK4 | 10 |

## Discussion

The availability of the genome sequences, decreasing sequencing cost, advances in mass spectrometry-based proteomics, access to open source data analysis pipelines and bioinformatics databases for biological interpretation have revolutionized all aspects of biological research in the post-genomics era. Advances made in understanding chicken health and disease are critical for the growth and sustainability of poultry production systems that play a crucial role in meeting the needs to feed the world’s increasing population. It is imperative to understand all aspects of the regulation of essential physiological processes to leverage the chicken genome sequence for gains in the US poultry industry. Regulatory cascades of given chicken pathophysiology include PTMs of proteins, which control multiple aspects of cell fate and are often deregulated in pathological conditions. Improving the existing annotation of chicken protein function is a critical step towards understanding the biological processes necessary for poultry health and disease prevention. Chemoproteomics enables enrichment of specific enzymes, despite their low abundance, by targeting active site. Therefore, chemoproteomics helps with the identification of low abundant enzymes that could remain undetected in an unfractionated proteome when analyzed by standard shotgun proteomics. Chemoproteomic approach has been applied to identify and quantify functional and active kinases, serine hydrolases, deubiquitinases, and other enzyme groups (27, 32–37). Phosphorylation, a reversible post-translational modification catalyzed by kinases acts as a molecular switch that regulates many signaling pathways that control several cellular processes including cell division, proliferation, differentiation, and apoptosis. Chemoproteomics-based identification of active kinases in chicken liver and spleen reported here is the first systematic description of this important class of enzymes in the chicken. Kinases reported in this study belong to AGC, CMGC, CAMK, CK1, STE, Tyrosine Kinase, Tyrosine Kinase-Like, RGC, and Atypical kinase groups. Majority of these kinases were previously uncharacterized in the chicken by genome-scale experimental approaches. Peptide arrays design includes short peptides sequences that are substrates for kinases and their design and application in poultry is focused on a subset of kinases involved in the regulation of immune and metabolic processes (23).

Peptide array based kinome studies in poultry focused on studying the host response to infection. Peptide arrays analysis of chicken macrophage kinome infected with different serovars of *Salmonella* identified differences in infection with two different strains (22), and evaluation of skeletal muscle kinome during Salmonella Typhimurium infection could identify metabolic changes that could alter fatty acid and glucose metabolism through AMPK and the insulin/mammalian target of rapamycin (mTOR) signaling pathways (38). A peptide array for studying the kinome of mallard and American black duck to study response to pathogens and environmental stress is available (39).

Pharmacological inhibition studies with piceatannol, a selective inhibitor of Syk tyrosine kinase, SB 203580, an inhibitor of p38 MAPK, U73122, an inhibitor of phospholipase C, and LY294002, an inhibitor of phosphoinositol-3 kinase established the role of these kinases in heterophil degranulation following the crosslinking of Fc receptors with IgG-bacteria during clearance of bacterial pathogens in poultry (40).

Pharmacological or biological inhibition of kinases in chicken can be alternatives to conventional antimicrobials. For example, SP600125, a specific inhibitor of JNK MAPK interferes with Fowl Adenoviruses (FAdVs) serotype 4 replication in Leghorn male hepatoma and enhances type I interferon production (41). FAdV-4 causes hydropericardium syndrome (HPS) and is associated with high mortality in fowl (41). Another application of specific kinase inhibition is to enhance bactericidal activity of macrophages using a protein kinase A inhibitor, H-89, that reversed the suppression of nitric oxide (NO) production during infection with *Salmonella* Enteritidis (42). Results of the present study to Identify catalytically active kinases in the chicken liver and tissue will complement ongoing efforts using peptide arrays and will contribute to understanding the role of this protein family in chicken development, growth and pathophysiology. Active Kinases from chicken liver and spleen will likely form the framework for interpreting future functional genomics studies in different domains of poultry research. Moreover, the identified kinome can be used as a starting point for chemoproteomic-based interrogation of kinase inhibitor selectivity. Kinases identified in this study are available through the Chickspress genome browser (*http://geneatlas.arl.arizona.edu/cgi-bin/GeneAtlas/chickspress_genes.cgii)*, for broader dissemination and biological discovery.

## Methods

### Biological material

The tissue samples were aseptically collected from commercial male broiler chickens at the end of their grow-out period (60-64 days). The birds were collected from the processing line immediately after they completed the stunning and exsanguination phase. Once removed, the intact bird carcasses we placed on ice and transported out of the processing plant for the collection of the tissue. Two birds were collected and processed for tissue removal to minimize the time from expiration to tissue removal. Once the tissues were aseptically collected, each sample was placed in a sterile whirl pack bag and placed on ice for transport to the laboratory.

Since the animal tissues were collected post-mortem, no IACUC was necessary for the work with animals. All methods were carried out in accordance with relevant guidelines and regulations, and all experimental protocols were approved by the Institutional Animal Care & Use Committee at the University of Florida (IACUC Study #201609318).

### Sample Preparation

ATP and ADP Acyl-nucleotide probes were synthesized as described previously (43). The sample preparation for mass spectrometry was done by ActivX Biosciences, Inc, USA. Chicken liver and chicken spleen tissues from two birds (two biological replicates) were lysed by sonication in lysis buffer (50 mM HEPES, pH 7.5, 150 mM NaCl, 0.1% Triton-X-100, phosphatase inhibitors [Cocktail II, AG Scientific]) by using the Omni Bead Ruptor 24. After lysis, the samples were cleared by centrifugation (16,200g, 15 min), and the supernatant collected for probe-labeling. Samples were incubated with 50 µL of a 10X aqueous solution of the desthiobiotin-adenosine triphosphate-acyl phosphate probe (ATP probe) or desthiobiotin-adenosine diphosphate-acyl phosphate probe (ADP probe) for a final probe concentration of 20 µM, for 10 min. Samples were prepared for MS analysis as described previously (24). Briefly, probe-labeled lysates were denatured and reduced (6 M urea, 10 mM DTT, 65°C, 15 min), alkylated (40 mM Iodoacetamide, 37°C, 30 min), and gel filtered (Biorad Econo-Pac 10G) into 10 mM ammonium bicarbonate, 2 M urea, 5 mM methionine. The desalted protein mixture was digested with trypsin (0.015 mg/mL) for 1 hr at 37°C, and desthiobiotinylated peptides captured using 12.5 μL high-capacity streptavidin resin (Thermo Scientific). Captured peptides were then washed extensively, and probe-labeled peptides eluted from the streptavidin beads using two 35-μL washes of a 50% CH_3_CN/water mixture containing 0.1% TFA at room temperature.

### LC-MS/MS Analysis

Samples were analyzed by LC-MS/MS as described previously (24). First, the samples were run in a data-dependent mode in ion trap instrument, and after the initial kinome was established, the identified ATP/ADP probe-labeled peptides were targeted based on the m/z and migration time by using the same ion trap mass spectrometer. Briefly, samples were analyzed on Thermo LTQ ion trap mass spectrometers coupled with Agilent 1100 series micro-HPLC systems with autosamplers, essentially as described (24, 44). HPLC grade solvents were used in all cases and were comprised of (A) 5% ACN, 0.1% formic acid, and (B) 100% ACN, 0.1% formic acid. Samples were loaded onto a pre-column peptide capTrap, desalted, and concentrated at 5% solvent B. Probe-labeled peptides were separated in 0.18 × 100 mm columns packed with 5 μm diameter, 300 Å Magic C18 stationary phase (Michrom Bioresources) equilibrated at 15% solvent B. Peptide samples were then separated using a three-stage linear gradient: 15-30% solvent B from 0-100 min.; 30-50% solvent B from 100-115 min.; 50-95 % solvent B from 115-120 min. The column flow rate was set to 2 μl/min. The nanospray source (Thermo Scientific) was operated with the spray voltage at 1.6 kV, the capillary temperature at 200°C, the capillary voltage at 46 V, tube lens voltage at 120 V, and relative collision energy set to 35%. Samples were first to run in data-dependent mode to identify probe-labeled peptides corresponding to kinases. The migration times and m/z values of these peptides were then assembled into a target list, and samples were subsequently run with this chicken kinase target list to allow for the relative quantitation of kinases in liver and spleen. The data were searched against the UniRef100 sequence database, taxonomy ID 9031, *Gallus*, by using the Sequest algorithm as described (24, 44).

For the quantitative analysis, integrated peak areas were determined by extracting the signal for theoretical fragments ions from each specifically targeted parent ion (corresponding to a known probe-labeled kinase peptide). The mass spectrometry proteomics data have been deposited to the ProteomeXchange Consortium via the PRIDE (45) partner repository with the dataset identifier PXD011548. Moreover, the proteomic data have been made available at the following link: *http://geneatlas.arl.arizona.edu/cgi-bin/GeneAtlas/chickspress_genes.cgii*

### Sequence Alignments

MS peptide results were mapped to *Gallus* proteins present in the UniProt database. *Gallus* proteins were selected for sequence alignment, and their sequence was blasted in NCBI to locate human and mouse orthologs, followed by alignment of these sequences.

### Chickspress Genome Browser

Chickspress genome browser is an open source resource that is a repository of proteomics, transcriptomics, miRNA and trait/QTL data from different tissues. Kinases identified in this study (liver and spleen) are made publicly available through this genome browser (*http://geneatlas.arl.arizona.edu/cgi-bin/GeneAtlas/chickspress_genes.cgii*). Unique identifications from the mass spectrometry data were searched against the UniProt database (as of April 2018) and current members for the Uniref clusters were downloaded inspected to ensure that they included chicken proteins. Chicken UniProt proteins were mapped a unique NCBI Gene records, and any discrepancies were reviewed manually. NCBI Gene records were used to map to the NCBI Galgal5 genome assembly, and the start and stop coordinates in the chromosome were retrieved and marked in the browser.

### Ingenuity Pathway Analysis

To identify the molecular functions, signaling pathways, toxicological functions, associated with the identified chicken kinases, we performed an Ingenuity Pathway Analysis (IPA) as described earlier (46). The IPA analysis is based on human, mouse and rat data not on chicken and therefore as a first step, human, mouse or rat orthologs of identified chicken proteins were identified. IPA mapped proteins from our dataset to gene objects (focus genes) in the Ingenuity Pathways Knowledgebase (IPKB) known to interact with other genes in networks and pathways. Fisher’s exact test was used to calculate the P-value to determine the probability of each biological function/disease or pathway being assigned by chance and a P ≤ 0.05 was used to select highly significant biological functions and signaling pathways.

### Immunoblotting

An equal amount of protein from each one of three liver or spleen samples (each tissue was harvested from a different bird) was loaded on a Criterion XT 4-12 % gradient polyacrylamide gel (Bio-Rad, USA) and proteins were separated by SDS-PAGE, followed by transfer onto Polyvinylidene difluoride (PVDF) membrane. A membrane was blocked with 5% non-fat dry milk in PBS containing 0.1% Tween (PBST) for 60 min at room temperature, and incubation with a primary anti-IKKa, AKT1 or PIK3CA rabbit antibodies (Cell Signaling Technology, USA) and goat anti-rabbit secondary HRP-conjugated antibody (Dako Scientific, USA). The antibodies recognized human proteins, which were over 95% identical to chicken proteins apart from IKKa, which was ∼79% identical, but the molecular weight of bands was compared to the predicted molecular weight of these proteins. All antibodies were diluted in Phosphate-buffered saline (PBS) containing 1% Tween-20 (PBS-T) and 1% non-fat dry milk. After extensive washes in PBS-T, Clarity Western Enhanced chemiluminescence (ECL) Substrate (Bio-Rad, USA) was applied, and the membrane was imaged by exposing signals onto an X-ray film.

## Acknowledgments

We would like to acknowledge Tyzoon Nomanbhoy and Brian Nordin for technical assistance with the generation of the chemoproteomics data. We would also like to thank Gary Jones for help with the analysis and thank Dr. Fiona McCarthy (University of Arizona) for her support with Chickspress. This study was supported by a Biotechnology Risk Assessment Grant Program No. 2016-31200-06012 from the National Institute of Food and Agriculture and Agricultural Research Service (NIFA).

## Author Contributions

MJE and BN jointly supervised the work. CRG and MO helped with data analysis, WWH, MO, BN, and MJE prepared figures, RB obtained tissue sections, MJE, BN, WWH and MJE wrote the manuscript.

